# Paleo-oscillomics: inferring aspects of Neanderthal language abilities from gene regulation of neural oscillations

**DOI:** 10.1101/167528

**Authors:** Elliot Murphy, Antonio Benítez-Burraco

**Affiliations:** Division of Psychology and Language Sciences, University College London, London, UK; Department of Psychology, University of Westminster, London, UK; Department of Spanish, Linguistics, and Theory of Literature. Faculty of Philology, University of Seville, Seville, Spain

**Keywords:** Language evolution, language disorders, Neanderthals, candidate-gene approach, neural oscillations, methylation patterns

## Abstract

Language seemingly evolved from changes in brain anatomy and wiring. We argue that language evolution can be better understood if particular changes in phasal and cross-frequency coupling properties of neural oscillations, resulting in core features of language, are considered. Because we cannot track the oscillatory activity of the brain from extinct hominins, we used our current understanding of the language oscillogenome (that is, the set of genes responsible for basic aspects of the oscillatory activity relevant for language) to infer some properties of the Neanderthal oscillome. We have found that several candidates for the language oscillogenome show differences in their methylation patterns between Neanderthals and humans. We argue that differences in their expression levels could be informative of differences in cognitive functions important for language.

## 1. Introduction

Language is the most distinctive human ability. Since it does not fossilize, language scientists instead need to rely on indirect evidence to gain some insight about how it evolved in our clade. Language is ultimately the product of neural activity and so skulls and brain endocasts are objects of particular interest. Because of the intimate cross-talk between the skull and the brain, we expect that the analysis of the hominin skull shape and growth pattern can inform how the brains of extinct hominin species were organized and how they developed during growth; factors which impact cognition (Zollikofer & Ponce de León, 2013). Language abilities in Neanderthals are a controversial issue (Johansson, 2015). The skull of anatomically-modern humans (AMH) differs both in size and shape from the skull of Neanderthals (Bruner, 2004). However, it is not still clear if calvarian differences correlate with endocast differences, and hence with brain function (Mounier et al., 2016), or if there is an AMH-specific brain developmental path (see Ponce de León et al., 2016; Williams & Cofran, 2016). Comparisons with extant primates are helping to refine our understanding of these differences (Pearce et al., 2013). Overall, gross differences in brain structure and function between hominin species have been used to infer differences in cognitive abilities important for language processing (Boeckx & Benítez-Burraco, 2014a).

In neighboring domains, neuronal oscillatory activity has been acknowledged as the basis of multiple cognitive and behavioural processes (Buzsáki, 2006). In our previous work we have argued that computational operations of language can be decomposed into generic processes that can be implemented via neural oscillations (Murphy, 2015a,b) and that certain multiplexing algorithms responsible for the construction of hierarchical phrase structures appear to be species-specific (Murphy, 2016a). We have also argued that neural oscillations provide a more reliable explanatory level for investigating linguistic computation than the (correlational) examination of nerve tracks and brain regions involved in language processing; not just in neurotypical language processing, but in cognitive conditions that are human-specific and entail language dysfunction (Benítez-Burraco & Murphy, 2016; Murphy & Benítez-Burraco, 2016). Meyer (2017) reviews a range of experimental evidence suggesting that oscillations play a causal role in language comprehension, from the chunking (δ) and storage (α) of phrasal units, to the prediction of upcoming syntactic material (β) and the unification of this material into a coherent semantic structure (γ). Beaudet (2017) points out that ‘[s]peech capacity cannot be appropriately inferred only from the cerebral condition, therefore hypotheses aiming at reconstructing the timing and mode of emergence of language in the hominin lineage should seek to combine various lines of evidence’ – with such lines of evidence including, we hope, links between genetics and oscillatory brain activity.

While we cannot track the oscillatory activity of extinct hominins, we believe that the next best solution to moving beyond this shortcoming consists in examining the coding regions and the expression patterns of the genes responsible for the brain’s oscillatory activity putatively responsible for human language. We have recently identified and functionally characterized a set of 48 genes that comprises the core of our *language oscillogenome*; that is, the set of genes responsible for basic aspects of the oscillatory activity relevant for language (Murphy & Benítez-Burraco, 2017). In this current contribution, we have looked for differences between Neanderthals and us in the coding regions of these genes, and in their methylation patterns, as well as signals of positive selection in our species. Our ultimate aim is inferring, from these genomic differences, differences in the brain activity important for language processing.

## 2. Evolutionary changes in the language oscillogenome

In order to achieve our objective, we first gathered via systematic literature review and database searches the available information concerning fixed changes in AMH proteins compared to Neanderthal proteins, genomic regions positively selected in AMHs after our split from Neanderthals, and differentially methylated regions (DMRs) in AMHs compared to Neanderthals. We have extensively built on Green et al’s (2010) seminal characterization of the Neanderthal genome, Prüfer et al.’s (2013) description of the Altai Neanderthal genome, and Gokhman et al.’s (2014, 2017) DNA methylation maps of modern humans and extinct hominins. We then looked for our candidates in these papers and databases. As noted in Murphy and Benítez-Burraco (2017), our candidates for the language oscillogenome fulfil three criteria: i) they are associated with language disorders (developmental dyslexia and/or specific language impairment) and/or language dysfunction in cognitive disorders entailing language deficits (schizophrenia and/or autism spectrum disorder), ii) they play a role in brain rhythmicity and/or are candidates for conditions entailing brain dysrhythmias, like epilepsy; and iii) gene-oscillations-language links can be confidently established for them.

Our findings are summarized in Table 1.

**Table 1.**
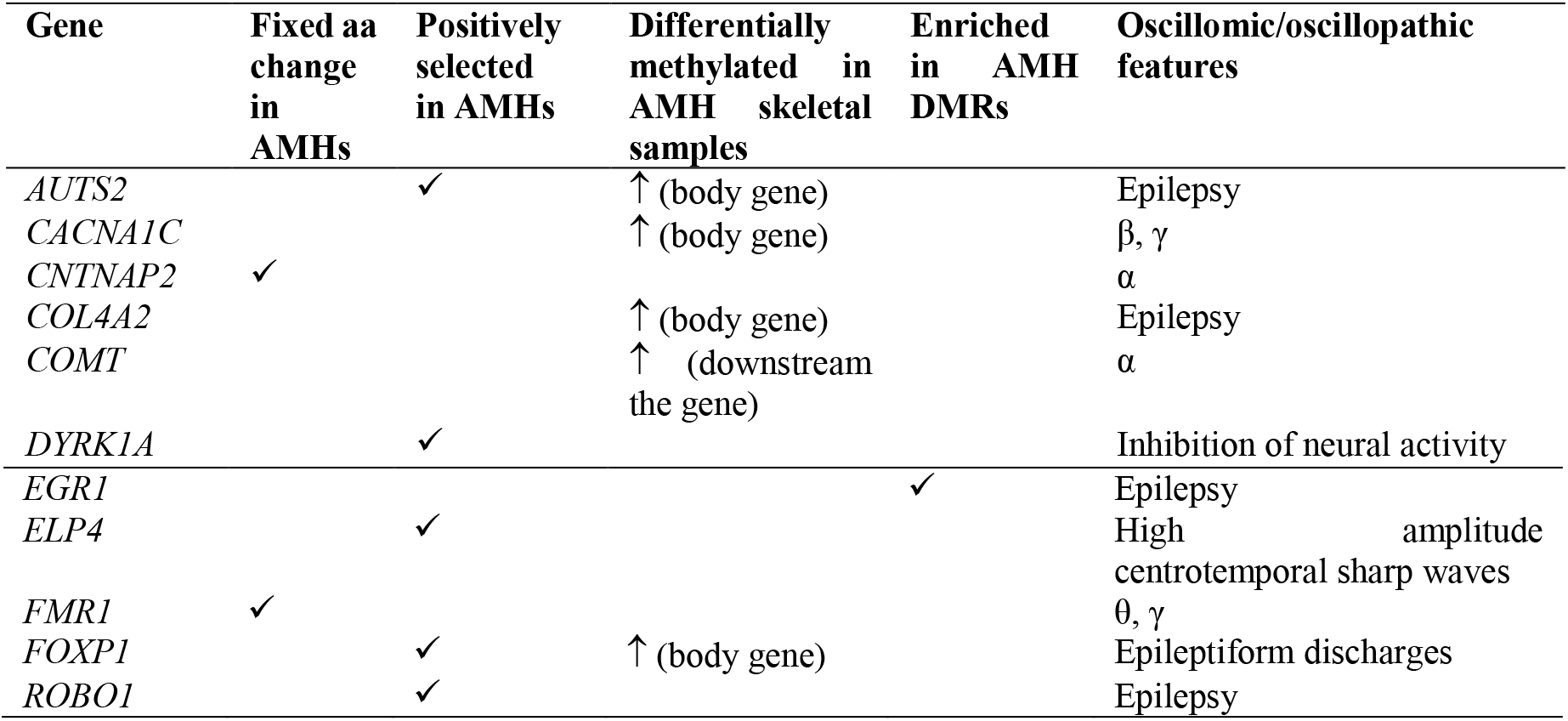
AMH-specific changes in genes involved in brain oscillations important for languages processing. The information about fixed aminoacid changes in the protein (column 1) derives for several sources, as noted in the main text. Evidence of positive selection (column 2) is from Green et al. (2010). The data about DMRs in AMH bone (column 3) are from Gokhman et al. 2017 (extended data table 2). The data about transcription factors enriched in AMH DMRs (column 4) is from Gokhman et al. 2014 (table S5). The information about the oscillomic/oscillopathic features of the gene is from Murphy and Benítez-Burraco 2017.

Among the 48 genes we highlighted in our previous work as part of the shared signature of abnormal brain oscillations associated with language deficits and core components of the language oscillogenome, 11 of these exhibit some difference between AMHs and Neanderthals. Two proteins bear fixed changes in one position in AMHs, although these changes are not expected to impact significantly on their structure and function: CNTNAP2 exhibits a Ile345Val change (reported as benign according to PolyPhen, and as tolerated (0.79) according to SIFT), whereas FMR1 shows a nearly fixed Ser145Ala change (frequency of 0.98 in AMHs) (classified as benign according to PolyPhen, and as tolerated (1) according to SIFT). More interestingly, 5 of our candidates are found within the regions that exhibit the strongest signals of positive selection in AMHs compared to Neanderthals, according to Green et al. (2010): *AUTS2, DYRK1A, ELP4, FOXP1*, and *ROBO1*. These positively selected regions contain numerous binding sites for transcription factors (Supplementary file, S1), which might have been affected by AMH-specific changes. Nonetheless, the most direct evidence of putative differences in the expression pattern of genes important for language under an oscillomic lens derive from the methylation maps of the AMH and the Neanderthal genomes obtained by Gokhman and colleagues (2014, 2017). We have found that several regions within the body gene of 4 of our candidates are hypermethylated in skeletal samples of AMHs compared to Neanderthals (*AUTS2, CACNA1C, COL4A2*, and *FOXP1*). Likewise, one hypermethylated region is found downstream one of our candidates (*COMT*) (Table 1 and supplementary file, S2). All these regions contain binding sites for several transcription factors of interest, like FOXA1, FOXA2, EP300, CBPB, MEF2A, FOXP2, FOXA2, CEBPB (Supplementary file, S2), which we have highlighted as important for language evolution, particularly, for changes related to the externalization of our linguistic thoughts (speech) (Boeckx & Benítez-Burraco, 2014b, Benítez-Burraco & Boeckx 2015). Finally, we have found that EGR1 is one of the enriched transcription factors in AMH DMRs.

Although methylation is commonly associated to promoter regulation, it is also a regulatory mechanism of gene expression when affecting to splice sites, coding regions, binding sites for transcription factors, and distal regulatory elements, like enhancers, silencers or insulators (Jones et al., 2012; Miller & Grant, 2013; Lev Maor et al., 2015). Accordingly, from the above differences in their methylation patterns, we might expect differences between Neanderthals and AMHs in the expression levels/patterns of several candidates for the language oscillogenome, namely, *AUTS2, CACNA1C, COL4A2, COMT, EGR1*, and *FOXP1*. Obviously, it is still to be determined whether the identified DMRs in skeletal methylation maps exist between the brains of AMHs and Neanderthals too. However, this might be the case (see Gokhman et al., 2016 for discussion).

Interestingly, duplications and deletions of all these genes, purportedly resulting in abnormal high or low levels of the protein, respectively, result in cognitive and language deficits (Table 2 Supplementary file, S4).

**Table 2.**
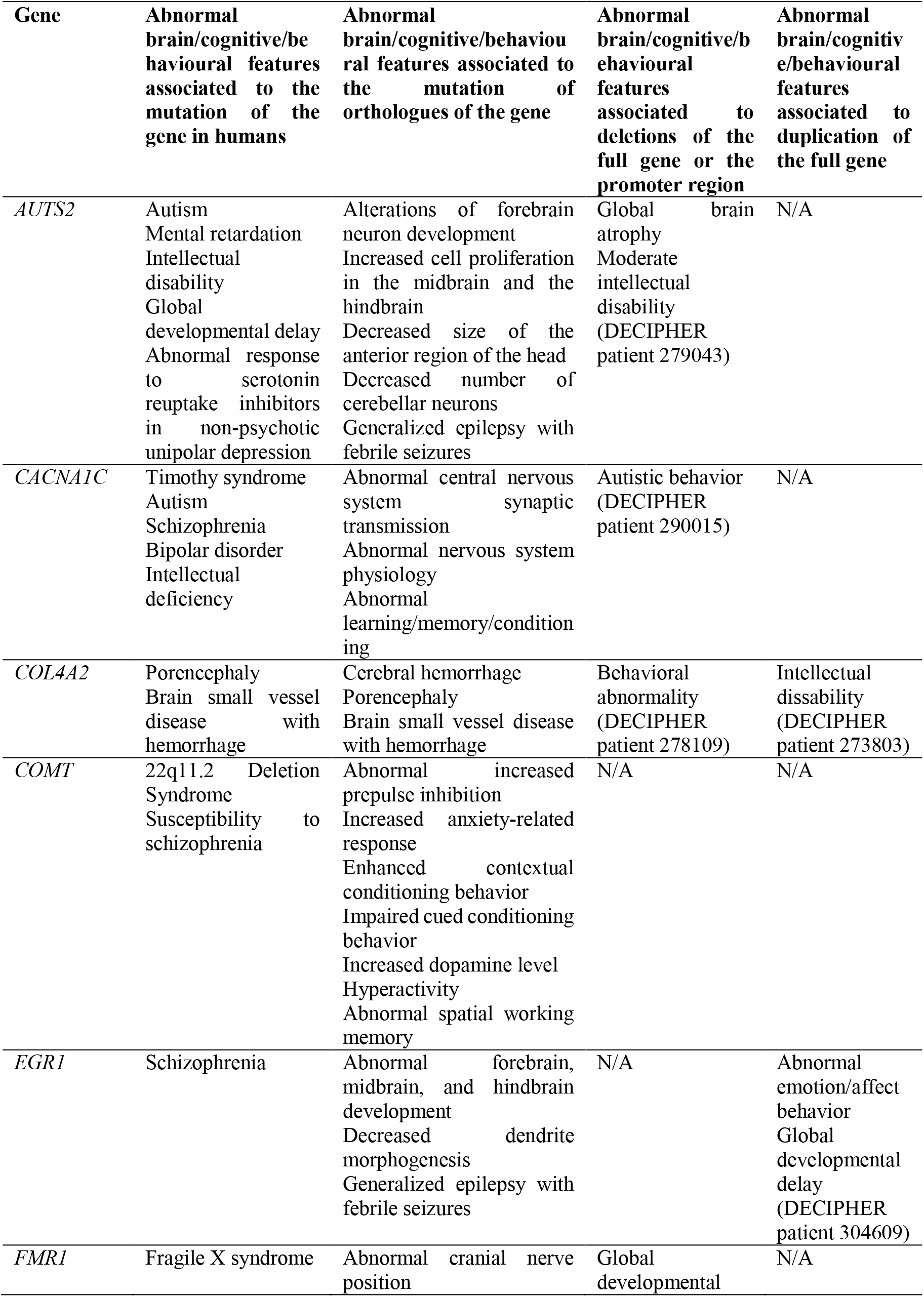
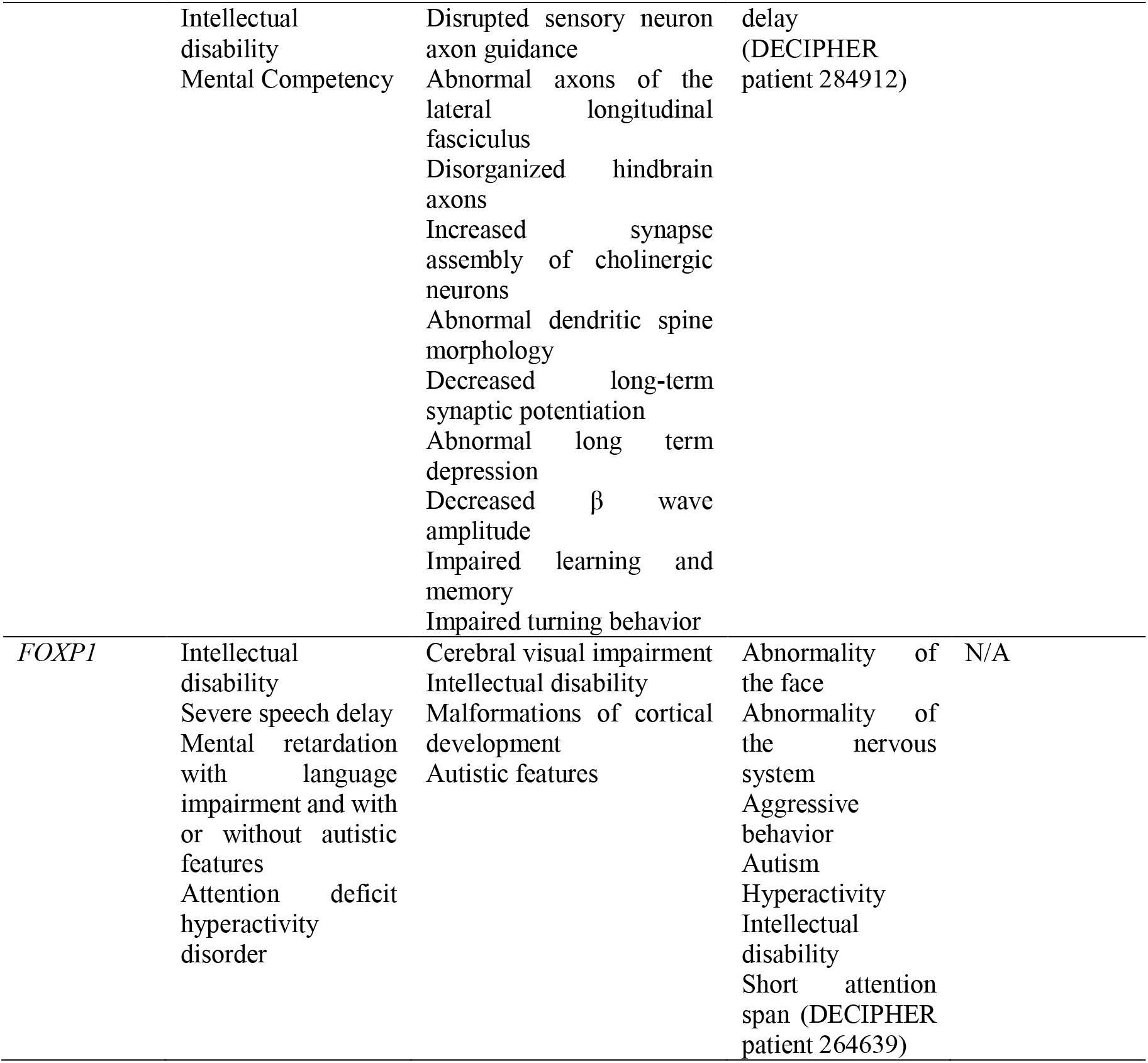
Abnormal brain, cognitive, and/or behavioural features associated to mutations in genes involved in brain oscillations important for languages processing. The abnormal features associated to the genes were retrieved from the Phenotypes tool of Ensembl (http://www.ensembl.org/index.html). Abnormal features observed in humans (column 1) derive from several databases of diseases and phenotypes directly associated with a gene, including the Online Mendelian Inheritance in Man (OMIM) compendium, Orphanet, and DDG2P, as well as from several sources of diseases and traits associated with variants of interest, including COSMIC, ClinVar, dbGaP, EGA, GIANT, HGMD-Public, MAGIC, NHGRI-EBI GWAS catalog, and UniProt. The abnormal phenotypes linked to mutations of the orthologe genes in other species come from several databases, including the International Mouse Phenotyping Consortium and Europhenome (for mouse), ZFIN (for zebrafish), Rat Genome Database (for rat), Animal_QTLdb and Online Mendelian Inhertiance in Animals. Because of biological differences between species, some animal phenotypes will not be easy to translate to our species. Columns 3 contains the available information about abnormal brain, cognitive, and/or behavioural features associated in humans to deletions of the full gene or the promoter region of the gene, as provided by DECIPHER (https://decipher.sanger.ac.uk/). Column 4 displays the abnormal brain, cognitive, and/or behavioural traits associated to duplications of the full sequence of the genes of interest, as also provided by DECIPHER.

*AUTS2* is involved in cytoskeletal regulation to control neuronal migration and neurite extension, but nuclear AUTS2 also controls transcription of several genes (Oksenberg et al., 2014; Hori & Hoshino, 2017). The gene has been associated with differential processing speeds (Luciano et al., 2011) and is a candidate for epilepsy (Mefford et al., 2010). *CACNA1C* encodes the alpha 1C subunit of the Cav1.2 voltage-dependent L-type calcium channel, involved in β and γ generation (Kumar et al., 2015). Pathogenic variants of *CACNA1C* have been associated with low performance scores in semantic verbal fluency tasks in schizophrenics (Krug et al., 2010), as well as to childhood-onset epilepsy, executive dysfunction, intellectual disability, and/or autism spectrum disorder (Damaj et al., 2015). *COL4A2* is a candidate for dyslexia, but it has been also associated to epilepsy and severe developmental delay (Giorgio et al. 2015, Smigiel et al. 2016). *COMT*, which encodes a catechol-O-methyltransferase that catalyzes the O-methylation of neurotransmitters like dopamine, epinephrine, and norepinephrine, has been regularly associated to schizophrenia, and particularly, to language dysfunction in this condition (Egan et al., 2001; Malhotra et al., 2002). The gene has been also associated to reading abilities in the neurotypical population (Landi et al., 2013), as well as to language performance and processing, and language acquisition, particularly to verbal fluency (Krug et al., 2009; Soeiro-De-Souza et al., 2013; Sugiura et al., 2017). Low activity levels in COMT has been associated to low voltage α (Enoch et al., 2003). Intriguingly, AMHs exhibit a hypomethylated region inside the body of one gene important for COMT function, namely *MSRA*, compared to Neanderthals (Gokhman et al., 2014, Table S2). EGR1 encodes a transcription factor that contributes to regulate neural plasticity and memory consolidation (Veyrac et al., 2014). In human epileptic foci, *EGR1* expression correlates with the frequency, amplitude and area of the interictal spikes, a hallmark of epileptic neocortex (Rakhade et al., 2007). Interestingly, *EGR1* is a target of FOXP2, the renowned “language gene” (Konopka et al., 2009), whereas EGR1 downregulates *PLAUR* (Matsunoshita et al., 2011), another of FOXP2’s targets (Roll et al., 2010). *FMRP1* is the main candidate for Fragile X syndrome (Kaufmann et al., 2004; Smith et al., 2012). *Fmr1*-knockout mice exhibit abnormal patterns of coupling between θ and γ oscillations in the CA1 area of the hippocampus, enhanced mGluR5 signalling resulting in altered neocortical rhythmic activity, and abnormally weak changes during tasks involving cognitive challenge (Hays et al., 2011; Radwan et al., 2016). FOXP1 interacts with FOXP2 to form functional heterodimers and *FOXP1* haplo-insufficiency has been associated with epileptiform discharges, speech delay, and motor delay (Carr et al., 2010).

Although in this paper we are interested in changes that are specific to AMHs and that allegedly contributed to the emergence of our species-specific pattern of brain oscillations, we wish to note that several of our candidates are differentially methylated in Neanderthals compared to other hominins. These differences might help account as well for their distinctive cognitive abilities, as discussed below. Accordingly, the body gene of *COMT* and *KANSL1* are differentially hypomethylated in bone samples of Neanderthals compared to AMHs. Likewise, one region downstream *ERG1* is found hypomethylated in Neanderthals (Supplementary file, S3). We have already discussed both *COMT* and *ERG1*. Regarding *KANSL1*, it encodes a component of the NSL1 complex, which functions as a regulator of gene transcription and chromatin organization, and is a candidate for Koolen-de Vries syndrome, which presents with epilepsy, developmental delay, and moderate intellectual disability, that mostly affects expressive language abilities (Koolen et al., 2016).

## 3. The language network in Neanderthals: Paleoneurology and oscillomics

Although very indirect due to the lack of actual evidence in the case of the former, comparative studies of Neanderthal and AMH cognition are vital if we are to hone our understanding of cognitive features unique to our species. As Mounier et al. (2016: 22) note, ‘the use of palaeoneurology to infer phylogenies of our genus is rare’. An emerging consensus indicates that while the skull size and shape of AMHs and Neanderthals are comparable, their internal neural organisation was in a number of respects distinct (Pearce et al., 2013). For instance, Neanderthals lived at much higher latitudes than AMHs and would have likely had larger eyes and, consequently, larger visual cortices (Pearce & Dunbar, 2012; Balzeau et al., 2012). Over the past couple of decades, digital anatomy and computed morphometrics have led to a seismic shift in functional craniology, exposing the generally plesiomorph structure of the Neanderthal braincase (Weber & Bookstein, 2011; Bruner, 2014; Williams & Cofran, 2016). Figure 1 summarises the major findings of this research. Naturally, not every neuroanatomical difference between Neanderthals and AMHs will be relevant to evolutionary linguistics, but we will conservatively review those differences which strike us as most significant. Notably, Neanderthals display upper parietal bulging which approximates the AMH shape but crucially lacks what Bruner (2014: 123) describes as ‘the overall bulging of the parietal surface characterizing the modern human brain globularity, in which the parietal proportions also enlarge in the longitudinal and vertical directions’. As we have already noted (Murphy & Benítez-Burraco, 2016), the globularity of the AMH braincase appears to have had direct consequences for the efficiency of oscillatory couplings across the cortex and subcortex which we have, in turn, used to derive the cross-modular nature of linguistic representations. Bruner (2014: 123) concludes that the type of parietal changes exhibited by the AMH brain likely resulted in modifications to visuo-spatial integration and ‘mental experiment capabilities’, which we interpret as recursively generated symbolic manipulations or what Tattersall (2017: 65) regards as ‘imagination’. Likewise, due to the expanded occipital lobe and visual cortices in Neanderthals, and due to the role of this region in α-inhibition vital to the coordination of cross-cortical feature binding (Murphy, 2016a), it is a possibility that Neanderthals were capable of executing a degree of control over representation integration not seen in other primates close to AMHs, but nevertheless still reduced in cross-cortical scope relative to AMHs. Further support for this hypothesis comes from Table 2: *COMT* exhibits hypermethylation in AMHs compared to Neanderthals, and since low-voltage α has been associated with low activity levels in the gene (Enoch et al., 2003), with *COMT* additionally being associated with language performance and verbal fluency (Krug et al., 2009; Soeiro-De-Souza et al., 2013), it is reasonable to assume that this hypermethylation contributes to the type of α-inhibition required to control inter-modular featural integration.

**Figure 1.**
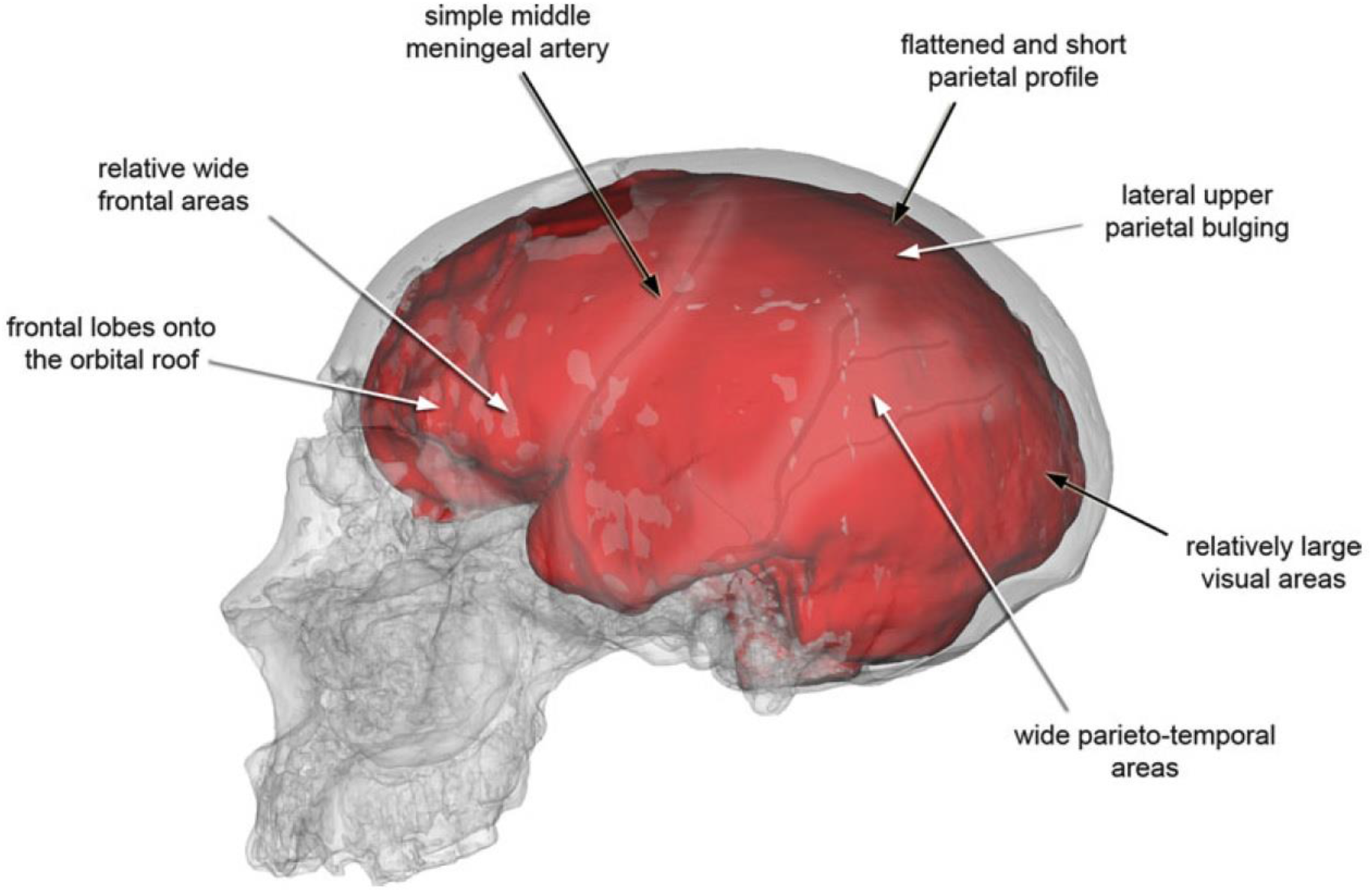
Paleoneurological characters of Neanderthals from a replica of the skull and endocasts of Saccopastore 1. White arrows indicate derived traits while black arrows indicate plesiomorph traits, which impose constraints as a result of large brain size (reproduced from Bruner 2014: 123).

It was shown in Murphy (2016b) that a range of monkeys (putty-nosed monkeys, Campbell’s monkeys, King Colobus monkeys, and New World monkeys) seem capable only of single-instance serial order concatenation and not hierarchical phrasal construction. Drawing on much neuroethological data, the likely oscillatory basis of this capacity was explored. It was proposed that phase-amplitude coupling between parahippocampal θ and cross-cortical γ was responsible for constructing individual call representations, and that primarily basal ganglia β increases signalled the maintenance of such calls in memory (Murphy, 2016b).

At this point, we need to distinguish between the possibility that Neanderthals were capable of speech from the possibility that they were capable of internally generating hierarchically organised sets of mental representations, as in natural language (as assumed in much work in generative linguistics; Chomsky et al. 2018). Neanderthals may have been capable of speech similar to AMHs due to the hard portion of the hyoid apparatus typically being preserved and their aural ability to be sensitive to frequencies (up to 5Hz) relevant for speech (Martinez *et al*., 2004). But what about the generation of hierarchically organised representations, the core of natural language (Chomsky et al. 2018)? We believe that since current work is revealing an oscillatory code for language, and since the molecular basis of this code is in turn becoming amenable to reconstruction (Murphy & Benítez-Burraco, 2017), the most reliable existing method of examining whether Neanderthals had anything resembling a proto-language is to draw inferences from emerging genetic databases. For instance, while Neanderthals may have been sensitive to language-relevant frequencies, the question we are interested in is whether neural assemblies internally oscillating at the same frequencies were used in the service of constructing hierarchically organised representations. Relatedly, Mithen (2014) reviews the evidence for cognitive similarities and differences in AMHs and Neanderthals and concludes, for example, that while pigment use likely indicates a complex social and emotional life, it does not provide evidence for symbolic thought.

It is likely that the human-specific oscillatory profile documented in Murphy (2016a), differing from the Neanderthal oscillome proposed here, is responsible for Mithen’s (1996, 2014) seminal distinction between the domain-specific Neanderthal mind (in which distinct core knowledge systems are largely isolated from each other) and the ‘cognitive fluidity’ of the AMH mind. Without this fluidity, integrating representations from distinct domains, Neanderthals ‘could never develop such [complex social] relationships and initiate a process of domestication’ (Mithen, 2014: 12; concerning domestication, see also Benítez-Burraco et al., 2016). More specifically, we would like to suggest that the distinct θ-γ code we predicted for the Neanderthal brain might also explain its presumably limited working memory (Wynn & Coolidge, 2012), given the crucial role that coupling between these rhythms appears to have in working memory operations across a number of modalities (Murphy, 2016a; Schomburg et al., 2014; Vosskuhl et al., 2015). A widening in the anterior fossa in Neanderthals (Figure 1) is associated with an expansion in Broca’s area, argued in Murphy (2016a) to be implicated in language-relevant memory buffers (see also Rogalsky et al., 2008 for a discussion of earlier proposals concerning Broca’s area in expanding working memory capacities). While linguistic representations may well have been assigned a special memory buffer (phonological loop) in Brodmann area 44, a part of Broca’s area (henceforth, BA 44), the computations associated with Merge, the core combinatorial operation in natural language that combines elementary linguistic units (Chomsky 1995), are certainly well distributed. In fact, even this perspective might not be wholly accurate: Along with BA 44 being a memory buffer (see Badre & Wagner 2007 for seminal discussion), it may also serve to select between competing syntactic interpretations, both storing and comparing representations (Novick et al. 2010). We confess that this perspective differs from the one articulated prominently by Friederici et al. (2017), who view parts of Broca’s area as playing a much more central role in phrase structure generation. Taken alongside the oscillogenomic evidence presented above – that Neanderthals may have exhibited reduced cross-frequency coupling between θ and γ due to differences in *CACNA1C* and *ELP4* expression – we should also conclude that the Neanderthal brain was likely incapable of exploiting this expanded Broca’s area, at least in the service of maintaining sets of symbolic representations. As with the partly globular parietal modifications discussed above, this appears to be another major feature of the Neanderthal brain which begins to approximate neuroanatomical characteristics that we hypothesize are required for core aspects of AMH syntax, but which ultimately displays other features which are likely incompatible with it. While the currently proposed oscillatory profile of Neanderthals does not necessarily exclude elements deemed necessary for phrasal construction (involving δ phase-amplitude coupling with θ and β, for instance; see Murphy, 2016b), the severe limitations which seem to have been imposed on it (most notably, working memory constraints and non-optimal cross-frequency couplings) suggests that even if Neanderthals did have core features of the neural code for AMH syntax there would have been a number of obstacles to implementing it.

Returning to Table 1, *CACNA1C*, which is differentially methylated in AMHs, contributes to β and γ generation, and has been correlated with semantic verbal fluency, may set AMHs apart from Neanderthals by contributing to speech-related fluency through β (implicated in storing ongoing speech representations in memory) and γ (involved in the initial construction of discourse representations, as discussed above). Relatedly, the abnormal dendritic spine morphology associated with *FMR1*, along with the decreased β power (Table 2), strengthens this notion.

With respect to the evolutionary implications, we conclude that the documented differences in the oscillatory profiles of humans and Neanderthals not only explain why we have language and Neanderthals likely did not, but they also shed some light on language-related cognitive differences, in particular concerning working memory limitations. We remain silent on the issue of the type of word-like representations Neanderthals may have possessed, and have focused purely on computational capacity, although it is worth noting that the evidence for Neanderthal symbolic thought at least suggests that it is their processing capabilities which distinguishes them from us. Our view is therefore somewhat similar to the mainstream generative position on language evolution (insofar as this particular topic goes, that is), although our views on neural reorganization in our lineage differ substantially from, for instance, Berwick and Chomsky’s (2016, 2017) or Friederici et al.’s (2017) positions (although the position articulated by these authors is also currently compatible with the data we present, given their focus on the computational novelty of Merge). Be that as it may, it should also be stressed that the current contribution constitutes highly speculative work, with the differences between Neanderthals and humans being a hotly debated and controversial topic. We hope at least to have opened up new avenues for exploring these differences, even if our present conclusions are soon revealed to be inaccurate and premature.

## 4. Conclusion

Overall, it is plausible that the oscillatory differences we have documented between AMHs and Neanderthals are connected to the major cultural and technological novelties which occurred towards the end of the Middle Pleistocene and which distinguish AMHs from our closest relatives. Expanding our molecular understanding of the Neanderthal brain can contribute to expanding the line of inquiry into what we have here called paleo-oscillomics. While it is hard to imagine anything in the fossil record shedding light on the *representational* basis of the Neanderthal brain (Berwick et al., 2013), reconstructing the Neanderthal oscillome can at least reveal aspects of its *computational* capacity.

## Acknowledgments

This work was supported by an Economic and Social Research Council scholarship (1474910) to EM, and by funds from the Spanish Ministry of Economy and Competitiveness (grant number FFI-2013-43823-P) to ABB. The authors wish express their gratitude to David Gokhman for sharing his data on methylation maps of Neanderthals and humans, and for kindly reading an earlier draft of the paper.

## Author contribution

ABB wrote Section 1-2. EM wrote Section 3-4.

